# MERS-CoV NSP16 necessary for IFN resistance and viral pathogenesis

**DOI:** 10.1101/173286

**Authors:** Vineet D. Menachery, Lisa E. Gralinski, Hugh D. Mitchell, Kenneth H. Dinnon, Sarah R. Leist, Boyd L. Yount, Rachel L. Graham, Eileen T. McAnarney, Kelly G. Stratton, Adam S. Cockrell, Kari Debbink, Amy C. Sims, Katrina M. Waters, Ralph S. Baric

## Abstract

Coronaviruses encode a mix of highly conserved and novel genes as well as genetic elements necessary for infection and pathogenesis, raising the possibility for common targets for attenuation and therapeutic design. In this study, we focus on the highly conserved nonstructural protein (NSP) 16, a viral 2’O methyl-transferase (MTase) that encodes critical functions in immune modulation and infection. Using reverse genetics, we disrupted a key motif in the conserved KDKE motif of MERS NSP16 (D130A) and evaluated the effect on viral infection and pathogenesis. While the absence of 2’O MTase activity had only marginal impact on propagation and replication in Vero cells, the MERS dNSP16 mutant demonstrated significant attenuation relative to control both in primary human airway cultures and *in vivo.* Further examination indicated the MERS dNSP16 mutant had a type I IFN based attenuation and was partially restored in the absence of IFIT molecules. Importantly, the robust attenuation permitted use of MERS dNSP16 as a live attenuated vaccine platform protecting from challenge with a mouse adapted MERS-CoV strain. These studies demonstrate the importance of the conserved 2’O MTase activity for CoV pathogenesis and highlight NSP16 as a conserved universal target for rapid live attenuated vaccine design in an expanding CoV outbreak setting.

**Significance:** Coronavirus emergence in both human and livestock represents a significant threat to global public health, as evidenced by the sudden emergence of SARS-CoV, MERS-CoV, PEDV and swine delta coronavirus in the 21^st^ century. These studies describe an approach that effectively targets the highly conserved 2’O methyl-transferase activity of coronaviruses for attenuation. With clear understanding of the IFN/IFIT based mechanism, NSP16 mutants provide a suitable target for a live attenuated vaccine platform as well as therapeutic development for both current and future emergent CoV strains. Importantly, other approaches targeting other conserved pan-coronavirus functions have not yet proven effective against MERS-CoV, illustrating the broad applicability of targeting viral 2’O MTase function across coronaviruses.

## Introduction

The emergence of Middle East Respiratory Syndrome Coronavirus (MERS-CoV) in 2012 represents the second severe coronavirus to strike the human population since the beginning of the 21^st^ century (1). Similar to its predecessor severe acute respiratory syndrome (SARS) CoV, MERS-CoV is characterized by severe lung infection and high mortality rates (2). Associated with elderly patients and nosocomial spread, MERS-CoV is likely harbored in camel populations with periodic reintroductions into humans, followed by periodic nosocomial outbreaks in hospital settings (3). Importantly, with the continued rate of globalization, MERS-CoV remains an ongoing threat for future outbreaks both in and outside the Middle East, as evidenced by the large outbreak in South Korea (4). Together, these factors highlight the importance of examining CoV pathogenesis and developing conserved therapeutic targets for treatment of current and future emergent strains.

Like all members of the CoV family, MERS-CoV maintains a balance of conserved and novel viral proteins within its genome (5). A member of the group 2C coronavirus family, a wealth of distinct accessory open reading frame (ORF) and non-structural proteins (NSPs) have already been established to have important roles in modulating the host immune response (6). Similarly, a number of highly conserved viral proteins for structure, replication, and fidelity are also maintained in the CoV backbone (7). Among these, MERS-CoV NSP16 provides a potent target for therapeutic development. A 2’O methyl-transferase, coronavirus NSP16 has been implicated in capping viral RNA and preventing its recognition by intracellular sensor MDA5 and antiviral effectors including IFIT family members (8). Generation of mutants within the NSP16 KDKE active site resulted in IFN mediated *in vitro* and *in vivo* attenuation of both mouse hepatitis virus and SARS-CoV (9, 10). As such, an approach targeting MERS-CoV NSP16 might be anticipated to result in attenuation and potentially provide a universal platform for CoV vaccines against future emergent strains.

Using reverse genetics to target residues in the highly conserved active site, we evaluated MERS-CoV infection outcomes in the context of an inactive NSP16 (dNSP16). Consistent with previous studies in SARS-CoV (10), the dNSP16 MERS-CoV mutant maintained no significant attenuation in terms of replication or the initial host immune response. However, both primary human airway epithelial cells and *in vivo* studies in a MERS-CoV mouse model demonstrated robust attenuation of dNSP16 mutant growth and pathogenesis. Notably, attenuation was both IFN and IFIT1 dependent providing a clear mechanism for attenuation. Importantly, the dNSP16 mutant also provided robust protection against a lethal MERS-CoV challenge and maintained attenuation in the mouse adapted backbone. Together, the results illustrate the broad conservation and necessity of NSP16 in CoV pathogenesis and highlight targeting this protein as a rapid response platform for future emergent CoV strains.

## Results

A combination of structural and biochemical approaches has established a critical role for CoV NSP16 in 2’O methyltransferase activity (**Fig. 1A**). Stabilized by interactions with NSP10 (orange), NSP16 relies on a highly conserved KDKE motif (red) to mediate its activity (11). Previous alteration of this motif in both group 2b SARS-CoV (10) and group 2a MHV (9) disrupted 2’O methyltransferase activity and attenuated varying aspects of infection. Based on high conservation across the CoV family (**Fig. 1B**), we hypothesized that disruption of the KDKE motif would also attenuate other emerging CoV families, including the group 2c MERS-CoV. Utilizing a MERS-CoV reversed genetic system (12), we disrupted the KDKE motif by mutating two nucleotides to produce a D130A change (**Fig. 1A**). The resulting disrupted NSP16 mutant (dNSP16) had no significant defect noted in stock titer generation (not shown); similarly, low MOI infection of both Vero cells and Calu3 cells, a respiratory epithelial cell line, demonstrated only modest attenuation at late time points (**Fig. 1C & D**). Together, these results indicate that NSP16 activity is not required for replication capacity.

**Figure 1.**
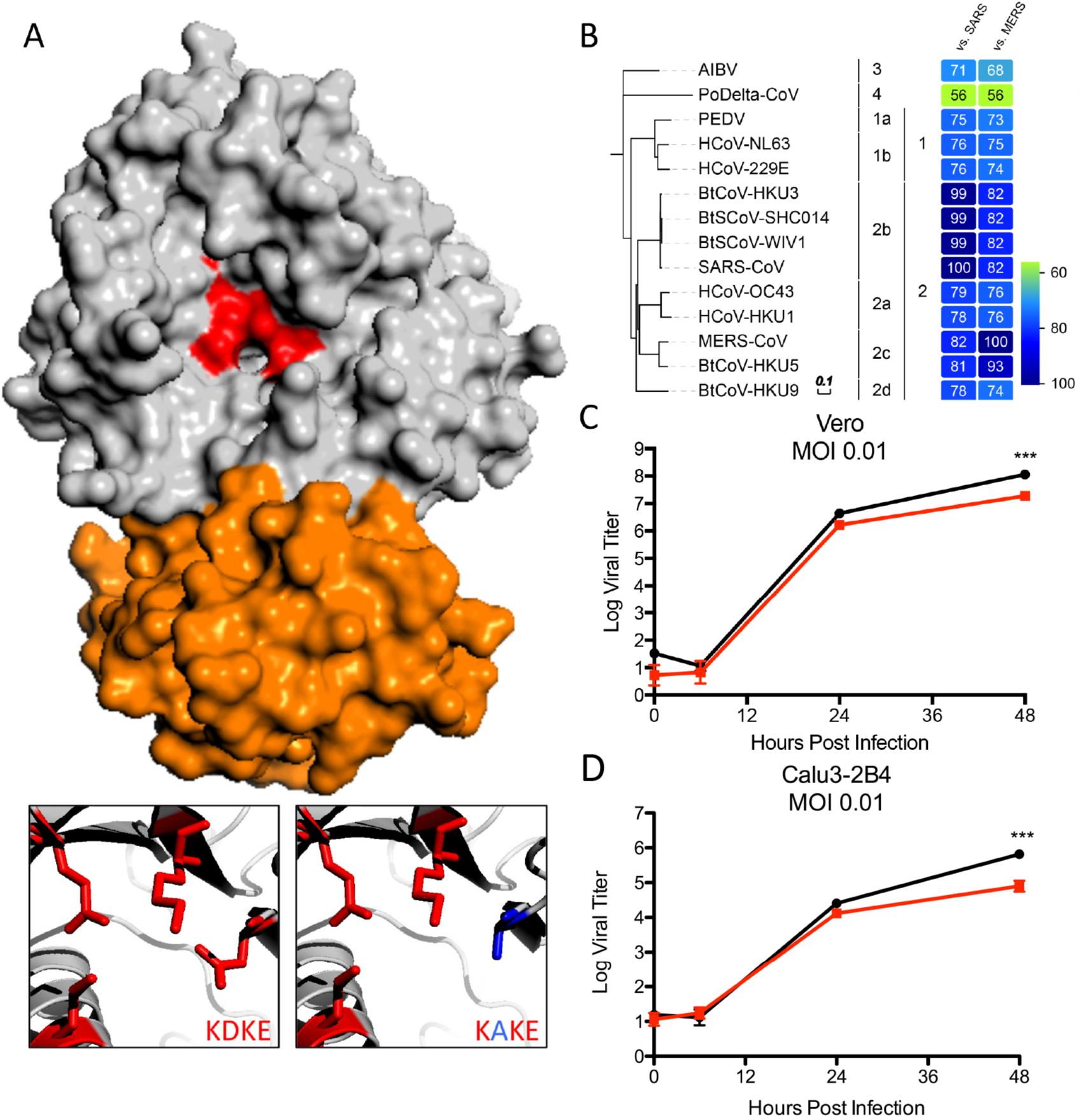
NSP16 highly conserved across coronavirus family. A) NSP16-NSP10 complex for MERS-CoV. NSP16 (gray) highlighting the conserved KDKE motif (red) required for 2’O methyl-transferase activity. Also shown, required NSP10 scaffold for MERS-CoV (orange). Inset displays conserved KDKE (right) as well as D130A (left) mutation that disrupts function. Homology models were created using Modeller in the Max-Planck Institute’s Bioinformatics Toolkit. The known crystal structure for the NSP10/16 complex (3R24 in the RCSB protein data bank) was used as the template structure (Chen et al., 2011). Homology models were then manipulated using MacPyMol. B) Heat maps were constructed from a set of representative coronaviruses from all four genogroups using alignment data paired with neighbor-joining phylogenetic trees built in Geneious (v.9.1.5) and visualized in EvolView (evolgenius.info). Trees show the degree of genetic similarity of NSP16 across CoV families. C & D) Viral replication of MERS dNSP16 mutant (red) relative to wild-type (WT) MERS-CoV (black) in (C) Vero cells and (D) Calu3 2B4 cells following MOI 0.01 infection. P-value representative of Student’s T-test with values representing ^⋆⋆⋆^ <0.001.

### *In vitro* host response similar between SARS and MERS dNSP16 mutants

Having established replication competence in both Vero and Calu3 cells, we next evaluated induction of host pathways following infection. Calu3 cells infected at MOI 5 demonstrated no differences in replication (not shown) and only modest differences in host induction (0 genes fold expression g >1.5 log2). Unlike previous studies with SARS-CoV, rapid cytopathic effect (CPE) by 24 hours limited analysis to early time points. Further DAVID based-analysis compared network host responses between MERS-CoV and SARS-CoV dNSP16 (**Fig. 2**). Over the first 24 hours of infection, both MERS-CoV and SARS-CoV dNSP16 showed no significant functional enrichment of any categories relative to corresponding wild-type (WT) infections, consistent with the lack of replication attenuation. However, at late times (>24 hours post infection), SARS-CoV produces robust changes in several host pathways including cytokine responses, inflammation, and extracellular activity. Similarly, changes in apoptosis, transcription repression, and regulation of phosphorylation indicated a host response more hostile to viral infection. While more rapid CPE following MERS-CoV dNSP16 infection precluded an equivalent finding at late time points, the SARS-CoV results suggest that the absence of NSP16 activity eventually initiates host response changes that contribute to attenuation at late time points.

**Figure 2.**
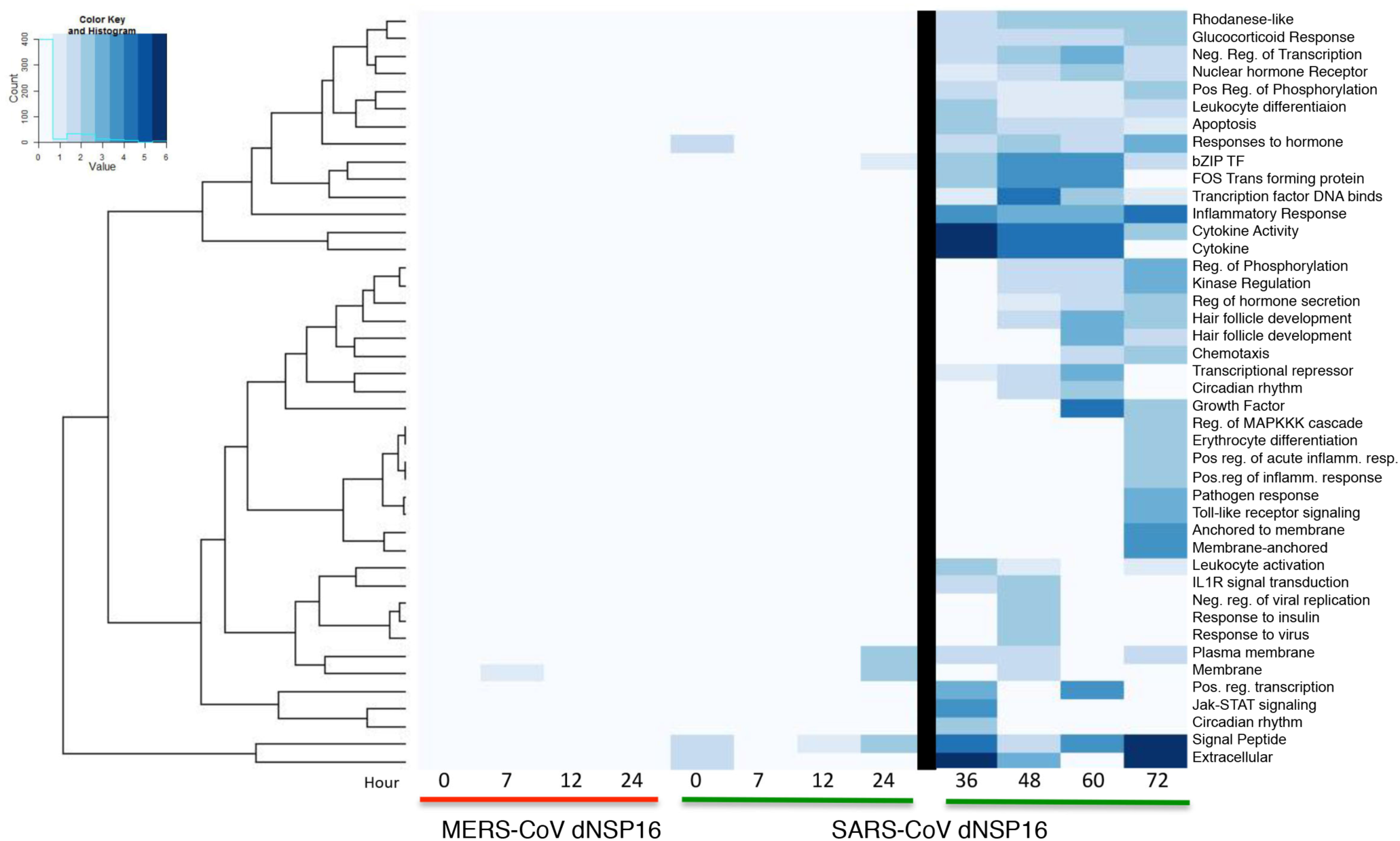
MERS dNSP16 infection produces minimal changes in early host responses. Changes in functional host gene clusters based on RNA expression following MOI 5 infection of Calu3 cells with MERS-CoV dNSP16 (left) or SARS-CoV dNSP16 (right) relative to wild-type (WT) control virus. Heat-map plots significant enrichment of clustered functional categories (as determined by DAVID analysis) for each mutant over time. Only marginal changes noted during the first 24 hours for both SARS and MERS-CoV dNSP16 mutants. After 24 hours (right) significant changes noted for SARS-CoV; MERS-CoV had significant cytopathic effect after 24 hours post infection precluding analysis.

### MERS-CoV dNSP16 attenuated in primary and *in vivo* models

To further examine the replicative capacity of dNSP16, we infected both human airway models and mice expressing human dipeptidyl peptidase 4 (DPP4), the receptor for MERS-CoV. Primary human airway cultures (HAE) were challenged with wild-type and dNSP16 MERS-CoV at a low MOI (**Fig. 3A**). While robust replication was observed following wild-type infection, dNSP16 MERS mutant had significant attenuation that corresponded well to previous results seen with SARS-CoV dNSP16 (10). We next examined MERS dNSP16 replication phenotypes in the context of *in vivo* infection using an adenovirus transduction model of BALB/c mice (13). While neither infection produced weight loss (not shown), WT MERS-CoV replicated efficiently at both day 2 and day 4 post infection (**Fig. 3B**); in contrast, no detectable replication was seen following infection with the MERS dNSP16 mutant. The lack of replication may be due to residual interferon responses associated with initial adenovirus infection. For greater clarity, we next infected CRISPR/CAS9 targeted mice that include mutations in mouse *Dpp4* at position 288 and 330 (288-330^+/+^) conferring efficient wild-type MERS-CoV infection and growth in mice, but no clinical disease (14). Following infection, no changes were observed in weight loss in either mice, consistent with previous findings (data not shown). However, absence of NSP16 activity severely attenuated dNSP16 replication at both day 2 and day 4 post infection (**Fig. 3C**). Coupled with data from HAE cultures and the adenovirus model, the results demonstrate clear attenuation of MERS dNSP16 relative to control.

**Figure 3.**
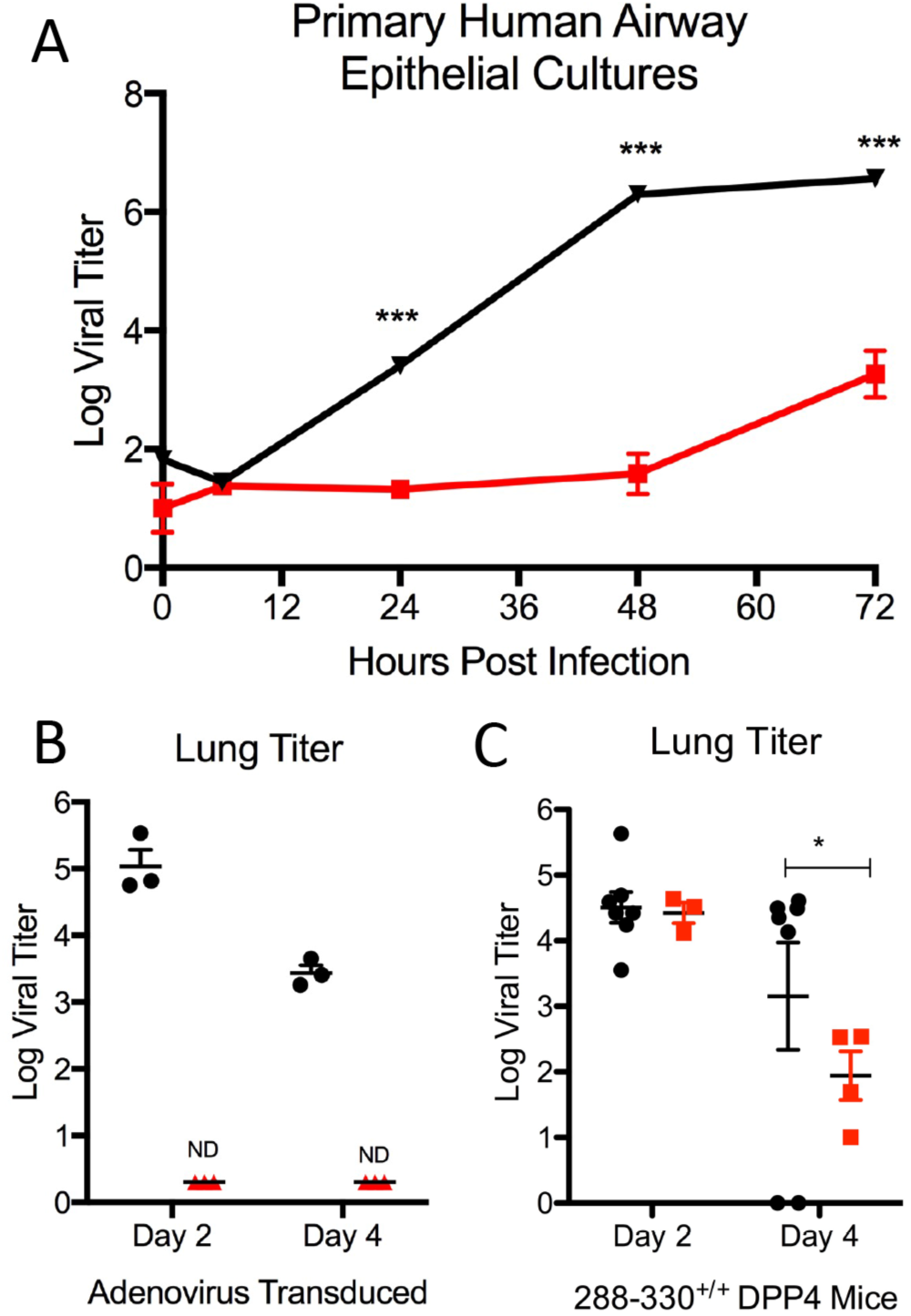
MERS dNSP16 attenuated in primary cultures and *in vivo.* A) Primary human airway epithelial cells infected with wild-type MERS-CoV (black) or dNSP16 mutant (red) at MOI 0.01 and monitored over time course. B) Day 2 and 4 lung titers from adenovirus transduced mice expressing human *DPP4* infected with wild-type MERS-CoV (black) or dNSP16 mutant (red). C) Day 2 and 4 lung titer from 288-330^+/+^ CRISPR mice infected with WT (Black) or dNSP16 (Red). P-value representative of Student’s T-test with values representing ^⋆^< 0.05 ^⋆⋆^ <0.01 ^⋆⋆⋆⋆^ <0.001.

### dNSP16 MERS-CoV attenuation mediated by IFN and IFIT1

Having established a deficit in MERS NSP16 mutant replication in relevant *in vitro* and *in vivo* models, we next sought to evaluate the mechanism of attenuation. Previous work by our lab and others had established increased susceptibility of NSP16 mutants to type I IFN (9, 10). While both viruses were sensitive to IFN treatment, the MERS dNSP16 mutant had a significant reduction in viral replication relative to control virus (**Fig. 4A**). These results are consistent with reports for NSP16 mutants in other coronaviruses (8). Extending this analysis further, we examined the role of *IFIT1* and *IFIT2* gene expression on this attenuation phenotype. Similar to SARS-CoV, knockdown of *IFIT1* augmented replication of the MERS dNSP16 mutant in the context of type I IFN pretreatment (**Fig. 4B**). In addition, knockdown augmented WT MERS-CoV infection, suggesting sensitivity to *IFIT1* activity despite the presence of NSP16. Notably, *IFIT2* knockdown had only a modest, non-significant impact on replication, contrasting results seen with SARS-CoV. Overall, the data indicates that MERS dNSP16 attenuation is driven by sensitivity to type I IFN mediated by the activity of *IFIT1*.

**Figure 4.**
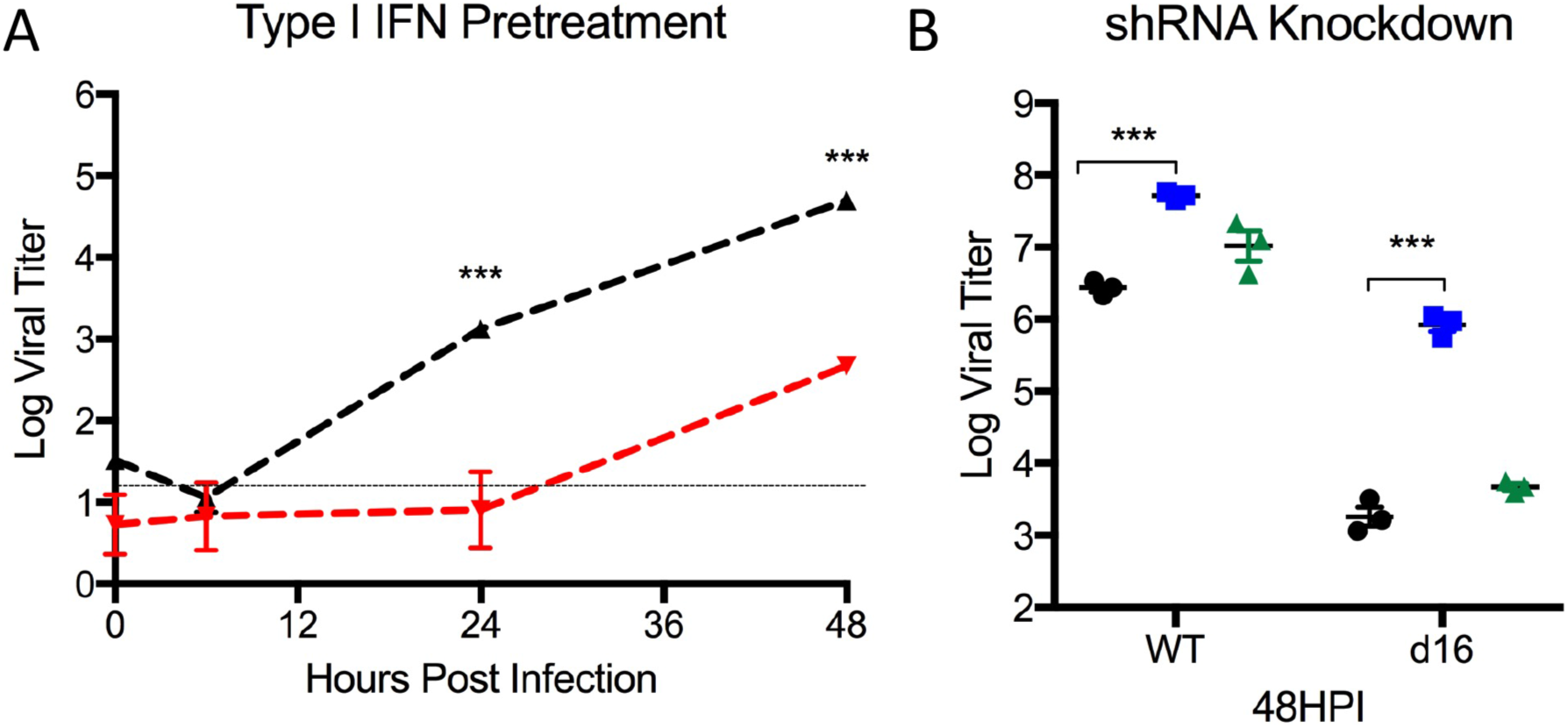
MERS dNSP16 attenuated by type IFN treatment via IFIT1. A) Vero cells treated with type I IFN (1000u) 16 hours prior to infection with either WT MERS-CoV (black) or dNSP16 mutant (Red). B) Vero cells expressing shRNA targeting *IFIT1* (blue), *IFIT2* (green), or no shRNA control (black) were pretreated with IFNβ (PBL Laboratories) and infected with MERS-CoV (left) or MERS dNSP16 (right). P-values based on Student T-test and are marked as indicated: ^⋆^<0.05, ^⋆⋆^<0.01, ^⋆⋆⋆^<0.001.

### NSP16 mutant vaccination protects from lethal MERS-CoV challenge

Based on IFN and IFIT1 attenuation phenotypes, targeting NSP16 offers a potential platform strategy for live attenuated vaccine generation. While previous work by our group had shown that the dNSP16 mutant of SARS-CoV conferred protection to lethal challenge, similar phenotypes in other more distant coronaviruses are essential for establishing universal principles of attenuation across a virus family. To test this hypothesis, DPP4 288/330^+/+^ mice were vaccinated with the wild-type MERS dNSP16 mutant and subsequently challenged with a mouse adapted MERS-CoV strain (**Fig. 5**) (14). Following challenge, dNSP16 vaccinated mice produced only modest weight loss, significantly contrasting severe disease seen in the control group (**Fig. 5A**). In addition, both viral replication and lung hemorrhage were significantly reduced in the context of the dNSP16 vaccine (**Fig. 5B & C**). Importantly, serum analysis revealed robust virus neutralization with values similar to sera from wild-type infected mice (**Fig. 5D**). Together, the results indicate that the dNSP16 MERS-CoV mutant can function as a vaccine platform that not only induces high levels of neutralizing antibodies, but provides compete protection from lethal MERS-CoV challenge.

**Figure 5.**
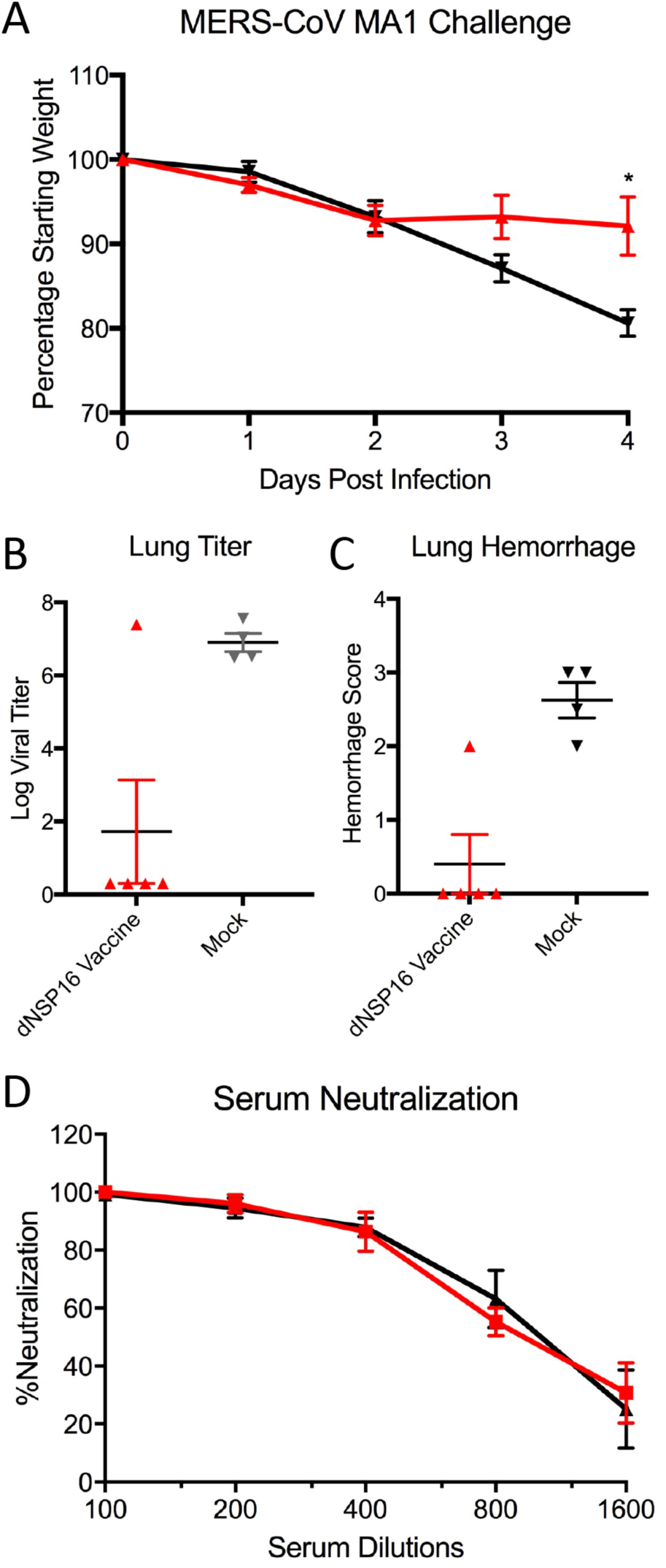
MERS dNSP16 mutants protects from lethal challenge. A) Weight loss, B) day 4 viral titer, and C) hemorrhage score following challenge of 288-330+/+ CRISPR/Cas9 mice vaccinated with wild-type MERS dNSP16 (red) or mock (black) with 10^^^6 pfu passaged mouse adapted MERS-CoV (14). D) Plaque reduction neutralization with sera from WT (black) or dNSP16 (red) vaccinated mice. P-value representative of Student’s T-test with values representing ^⋆^< 0.05 ^⋆⋆^ <0.01 ^⋆⋆⋆^ <0.001.

### NSP16 mutation attenuates mouse adapted MERS-CoV

Despite conferring protection in the wild-type MERS-CoV backbone, it was unclear if the NSP16 mutant would be sufficiently attenuated in a virulent MERS backbone. To address this question, we inserted the dNSP16 mutation (D130A) into the mouse adapted MERS-CoV backbone (14). Following infection, mouse adapted icMERS-CoV produced rapid weight loss and lethality (**Fig. 6A**). In contrast, the mouse adapted dNSP16 mutant had only modest weight loss and 100% survival following infection. In addition, viral replication of the dNSP16 mutant was significantly attenuated relative to WT at days 2 and 4 post infection (**Fig. 6B**). Finally, hemorrhage scoring of the lung revealed minimal disease in dNSP16 relative to control mice at day 4 post infection (**Fig. 6C**). Overall, the results demonstrate robust attenuation of MERS-CoV pathogenesis in the context a NSP16 mutation.

**Figure 6.**
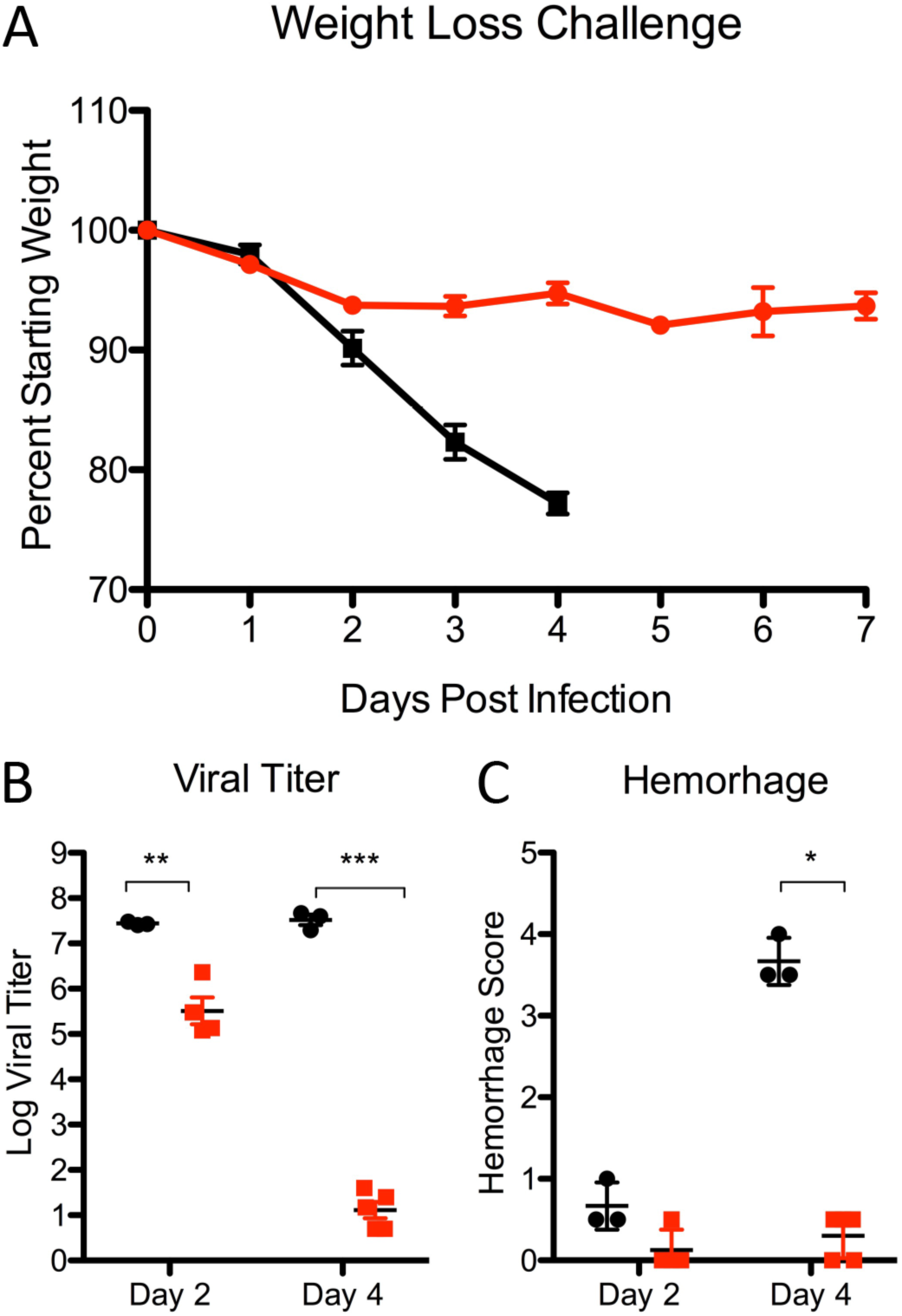
MERS dNSP16 mutant attenuated in virulent mouse adapted MERS-CoV strain. A) Weight loss, B) lung titers, and C) hemorrhage score following infection of 288-330^+/+^ CRISPR/Cas9 mice infected with 10^6 pfu MERS-CoV MA1 (black) or dNSP16 MA1 (red) at day 2 and 4. C) P-value representative of Student’s T-test with values representing ^⋆^< 0.05 ^⋆⋆^ <0.01 ^⋆⋆⋆^ <0.001.

## Discussion

In the context of the ongoing MERS-CoV outbreak, the development of universal platform strategies to attenuate emerging and contemporary coronaviruses is a significant priority. In this study, we demonstrate the critical importance of NSP16 function to MERS-CoV pathogenesis *in vitro* and *in vivo.* Similar to MHV and SARS-CoV (9, 10), the disruption of 2’O MTase activity in MERS-CoV had no significant impact on replication or early host response patterns *in vitro.* However, the dNSP16 MERS mutant demonstrated attenuated replication and growth in primary human airway cultures as well as reduced disease *in vivo*, relative to WT virus. Further examination revealed increased sensitivity to type I IFN in an IFIT dependent manner. Notably, this IFN/IFIT based attenuation phenotype provided robust protection from a lethal challenge following vaccination with the dNSP16 mutant. Importantly, the NSP16 mutant also was fully attenuated in a highly virulent mouse-adapted MERS-backbone. These results, coupled with previous work in other CoVs, highlight the viral 2’O MTase activity as a potential universal platform for therapeutic treatment and vaccine development for current and future emergent human and animal CoVs.

The attenuation of the MERS-CoV NSP16 mutant also provides evidence for targeting viral 2’O MTase as a global therapeutic strategy against emergent RNA viruses. Previous work in both flaviviruses and coronaviruses has demonstrated robust attenuation by targeting the conserved KDKE motif of the viral 2’O MTase (8, 15, 16). Importantly, 2’O MTase activity is the only known function for NSP16 and it appears completely dispensable for CoV replication in the absence of a strong IFN response (8). In contrast, flaviviruses without nsp5 encoded MTase activity are much more sensitive to innate immune responses (17); mutants in West Nile virus, Dengue, and Japanese encephalitis virus are highly replication attenuated at early time points (18, 19). While diminished virus yields likely reduces flaviruses vaccine utility *in vivo*, robust propagation of both MERS and SARS-CoV dNSP16 mutants provide a useful parameter for vaccine generation. For both viral families, drugs targeting 2’O MTase activity may have great efficacy if paired with stimulation of the interferon responsive genes including IFIT1 (8, 15). Overall, the results indicate that viral 2’O MTase activity is a critical viral determinant for pathogenesis and can be leveraged to develop therapeutic treatments.

NSP16 also represents the third highly conserved CoV protein targeted as the basis of a vaccine platform. Previous work by our group demonstrated the importance of CoV fidelity on infection and pathogenesis (20); disrupting the exonuclease activity of NSP14 rendered an attenuated, reversion proof virus in both MHV and SARS-CoV (20-22). For SARS-CoV, the disruption served as the basis of a successful live attenuated vaccine (20). Similarly, groups have targeted the envelope protein of SARS-CoV and defined a key inflammatory role for E protein in pathogenesis. SARS mutants targeting E function also conferred protection from lethal challenge (23, 24). For both NSP14 and E mutants, fidelity and inflammation induction are partially responsible for attenuation. However, both viral mutants also maintain replication attenuated in terms of kinetics and yields, potentially complicating their use as a vaccine platforms, especially in outbred populations (20, 24). Importantly, the NSP14 mutant has not yet been recovered within the context of MERS-CoV and failure has been reported in group I coronaviruses (25). Similarly, disruption of the E protein renders MERS-CoV, TGEV and MHV replication deficient without exogenous complementation (26-28); this broad attenuation of the delta E mutant potentially limits its use as a live attenuated vaccine. In contrast, MERS dNSP16 mutant maintains robust propagation as well as IFN/IFIT based attenuation phenotypes. Together, these results suggest that targeting NSP16 may be the most broadly applicable platform for CoV attenuation.

Despite the success of NSP16 mutants in protection studies with SARS and MERS-CoV, several additional parameters must be considered in the context of coronavirus vaccination. Previous work by our group has demonstrated failure of CoV subunit vaccines in aged animals and in the context of heterologous challenge (29). With the aged representing a population with high mortality and marginal vaccination success, efficacy of the NSP16 mutant in this population is paramount in its pursuit as a platform. Similarly, the existence of numerous CoVs in animal populations raises the concerns for heterologous challenge from emergent viruses (30, 31). Prior reports had also demonstrated vaccine induced disease following heterologous challenge with related SARS-like viruses (29, 32). With this in mind, NSP16 vaccinated mice will need to be examined in the context of heterologous challenge to determine if vaccine induced pathology occurs. Together, these two factors represent important checkpoints in the pursuit of NSP16 as a universal CoV vaccine platform.

In addition to aging and heterologous challenge, reversion and baseline pathogenesis also represent important risks that must be evaluated in the context of an NSP16 vaccine. While the NSP14 SARS vaccine was absent sterilizing immunity in immunodeficient mice, the lack of reversion over time *in vivo* indicates safety in the approach (20). In contrast, passage of the E mutant rendered a novel mutant that transplanted a critical ion channel function from another viral protein and thus restoring partial virulence (33). Studies examining reversion potential of NSP16 are critical prior to use as a vaccine; these concerns are especially important considering both the SARS and MERS dNSP16 mutants have robust replications permitting additional opportunities for reversion (10). In addition, equivalent host immune responses during the early infection may produce substantial damage prior to IFN/IFIT based attenuations, most notably in aged and immunocompromised mice. Therefore, despite the promise of successful attenuation in multiple CoV backbones, several additional metrics must be examined prior to using NSP16 mutants as a universal CoV vaccine platform.

Overall, the current study demonstrates that targeting 2’O MTase activity is a robust strategy to attenuate MERS-CoV and other emergent coronaviruses. In the absence of NSP16 activity, the MERS-CoV mutant is sensitive to type I IFN in an IFIT dependent manner providing clear mechanism for attenuation. Importantly, unlike other conserved CoV platforms, the NSP16 mutant is both viable and robust enough to be utilized as an effective live-attenuated vaccine. While further vaccine characterization is required, the results indicate that disruption of CoV NSP16 activity can be the basis for therapeutic strategies for both current and future emergent CoV infection both in human and animal populations.

## Methods & Materials

### Cells & Viruses

Wild-type, mutant, and mouse adapted MERS-CoV were previously described (12, 14) and cultured on Vero 81 cells, grown in DMEM or Optimem (Gibco, CA) and 5% Fetal Clone Serum (Hyclone, South Logan, UT) along with antibiotic/antimycotic (Gibco, Carlsbad, CA). Growth curves in Vero, Calu-3 2B4, and human airway epithelial cells were performed as previously described (9, 31). Briefly, cells were washed with PBS, and inoculated with virus or mock diluted in PBS for 40 minutes at 37 °C. Following inoculation, cells were washed 3 times, and fresh media added to signify time 0 hr. Samples were harvested at the described time points. For IFN pretreatments, 100 units/mL of recombinant human IFNβ (PBL Laboratories) was added to cells 16 hours prior to inoculation and infected as described above. Stable shRNA knockdown Vero cell lines for IFIT1 and IFIT2 were previously described (10). All virus cultivation was performed in a BSL3 laboratory with redundant fans in Biosafety Cabinets as described previously by our group (32, 33). All personnel wore Powdered Air Purifying Respirator (3M breathe easy) with Tyvek suits, aprons, booties and were double-gloved.

### Construction of Wild-type and Mutant NSP16 Viruses

Both wild-type and mutant viruses were derived from either MERS-CoV EMC or corresponding mouse adapted (MA1) infectious clone as previously described (12). For NSP16 mutant construction, the D130A mutation changed the sequence from MERS E fragment cloned within the pSMART vector (Lucigen) was used for Alanine scanning mutagenesis of conserved residues in nsp16. For the D130A change, a product was generated by PCR using primers against MERS-CoV NSP16. Fragment 1: EMC:E#2(+): TGAACTACCTGTAGCTGTAG EMC:EmuC(-): NNNNNN<u>GCTCTTCT</u>CGCGGAAATAACAAGATCCACTTG; Fragment 2: EMC:EmuC(+): NNNNNN<u>GCTCTTCC</u>GCGATGTATGATCCTACTACTAAG EMC:E#6(-): CAACCTCAATACAAGCAGAC. The two resulting products were digested with SapI (underlined) and ligated overnight with T4 DNA ligase. This product was then digested with PpuMI and NsiI and used to replace the region of the EMC E plasmid (puc57) which had been similarly digested. Thereafter, plasmids containing wild-type and mutant MERS-CoV genome fragments were amplified, excised, ligated, and purified. *In vitro* transcription reactions were then preformed to synthesize full-length genomic RNA, which was transfected into Vero E6 cells. The media from transfected cells were harvested and served as seed stocks for subsequent experiments. Viral mutants were confirmed by sequence analysis prior to use. Synthetic construction of mutants of NSP16 were approved by the University of North Carolina Institutional Biosafety Committee.

### RNA isolation, microarray processing and identification of differential expression

RNA isolation and microarray processing, quality control, and normalization from Calu-3 cells was carried out as previously described (34). Differential expression (DE) was determined by comparing virus-infected replicates to time-matched mock replicates. Criteria for DE in determining the consensus ISG list were an absolute Log_2_ fold change of >1.5 and a false discovery rate (FDR)-adjusted *P* value of <0.05 for a given time point.

### Clustering and Functional Enrichment

Genes identified as differentially expressed were used to generate clustered expression heat maps. Hierarchical clustering (using Euclidean distance and complete linkage clustering) was used to cluster gene expression according to behavior across experimental conditions. The DAVID online resource (david.ncifcrf.gov) was used to acquire functional enrichment results for the genes in each cluster. DAVID output was manually summarized for each cluster. Plots were generated with R.

### Ethics Statement

This study was carried out in accordance with the recommendations for care and use of animals by the Office of Laboratory Animal Welfare (OLAW), National Institutes of Health. The Institutional Animal Care and Use Committee (IACUC) of The University of North Carolina at Chapel Hill (UNC, Permit Number A-3410-01) approved the animal study protocol (IACUC Protocol #15-009 and 13-072) followed in this manuscript.

### Mouse Infections and Vaccinations

10-20 week old BALB/c (Envigo/Harlan) or CRISPR/Cas9 288-330^+/+^ C57BL/6 mice were anaesthetized with ketamine and Xylazine (as per IACUC, UNC guidelines) and intranasally inoculated with a 50 μl volume containing 10^6^ plaque forming units (pfu) of MERS-CoV WT, MERS-CoV dNSP16 virus, mouse adapted variants, or PBS mock as indicated in the figure legends. Infected animals were monitored for weight loss, morbidity, clinical signs of disease, and lung titers were determined as described previously (37). *In vivo* adenovirus transduction with *DPP4* were performed as previously described (13). For vaccination experiments, 10-20 week old 288-330^+/+^ mice were infected with 10^^^6 pfu of MERS dNSP16 as described above, monitored for clinical symptoms for 7 days, and then challenged 4 weeks post vaccination with 10^^^6 pfu MERS-CoV MA1 passaged virus. Animal housing, care and experimental protocols were in accordance with University of North Carolina (UNC) Institutional Animal Care and Use Committee (IACUC) guidelines.

### Data Dissemination

Raw microarray data for these studies were deposited in publicly available databases in the National Center for Biotechnology Information’s (NCBI) Gene Expression Omnibus (35) and are accessible through GEO Series: GSE65574. (http://www.ncbi.nlm.nih.gov/geo/query/acc.cgi?acc=GSE65574).

## Acknowledgements

Research was supported by grants from NIAID of the NIH (U19AI100625 to RSB; U19AI106772 to RSB; HHSN272201000019I-HHSN27200003 to RSB; K99AG049092 to VDM). Support for primary human airway cultures supported by NIH through UNC Cystic Fibrosis Research and Translation Core Center Cell Models Core at UNC (NIH P30DK065988). The content is solely the responsibility of the authors and does not necessarily represent the official views of the NIH. PNNL is operated by Battelle Memorial Institute for the DOE under contract number DE-AC05-76RLO1830.

